# WebMeV: a Cloud Platform for Analyzing and Visualizing Cancer Genomic Data

**DOI:** 10.1101/147884

**Authors:** Yaoyu E. Wang, Lev Kuznetsov, Antony Partensky, Jalil Farid, John Quackenbush

**Affiliations:** Center for Cancer Computational Biology; Dana-Farber Cancer Institute, Boston, MA

## Abstract

Although large, complex genomic data sets are increasingly easy to generate, and the number of publicly available data sets in cancer and other diseases is rapidly growing, the lack of intuitive, easy to use analysis tools has remained a barrier to the effective use of such data. WebMeV (https://mev.tm4.org) is an open-source, web-based tool that gives users access to sophisticated tools for analysis of RNA-Seq and other data in an interface designed to democratize data access. WebMeV combines cloud-based technologies with a simple user interface to allow users to access large public data sets such as that from The Cancer Genome Atlas (TCGA) or to upload their own. The interface allows users to visualize data and to apply advanced data mining analysis methods to explore the data and draw biologically meaningful conclusions. We provide an overview of WebMeV and demonstrate two simple use cases that illustrate the value of putting data analysis in the hands of those looking to explore the underlying biology of the systems being studied.

## Introduction

As the cost of sequencing has fallen, the number of large, publicly available genomic data sets has grown dramatically. Projects such as The Cancer Genome Atlas (TCGA) (1), the Genotype-Tissue Expression Project (GTEx) (2), and others have placed large, complex, multi-omic data into the public domain. These data sets provide unprecedented opportunities for users to perform complex analyses and re-analyses, to make new discoveries, to search for associations among genes and phenotypes, and to test hypotheses that can then be more fully explored in future experiments. However, technical challenges such as moving and analyzing large multi-omic data sets, and the lack of intuitive and easy to use tools for data analysis, have limited broad exploration of the available data, often preventing experimental biologists from directly interacting with the data. WebMeV (https://mev.tm4.org) is an open-source, web-based application designed to take advantage of cloud computing resources to provide users with access to cutting edge genomic analysis tools, direct access to large public domain data sets, intuitive user interfaces and data visualization, and tools for reproducible research.

## Approach

The WebMeV architecture is built around an application server that communicates with a computing server deployed on Amazon Web Services (AWS) to provide access to expandable computational power that takes advantage of modern cloud architectures. WebMeV was developed to provide an intuitive, user-friendly interface both large public data sets and the advanced analytical methods that have been released through Bioconductor (3), which is the largest publicly available collection of open source genomic data analysis tools.

### Bioconductor

Bioconductor is based primarily on the R programming language with analysis tools distributed as R packages. While Bioconductor receives contributions from an extensive community of bioinformatics and biostatistics developers, the majority of Bioconductor packages have only command line interfaces. Further, both Bioconductor and R require appropriate installation, configuration and version control to assure the analytical tools function and provide reproducible results. These factors present significant barriers for many potential users, including basic and translational researchers who are interest ed in analyzing and exploring complex genomic data—and who are often in the best position to interpret the results. So despite the considerable resource that Bioconductor represents, adoption by non-computational scientists has been slow. WebMeV was designed to lower those barriers by placing data and appropriate Bioconductor tools into a easy to use interface.

WebMeV uses Rserve (https://cran.r-project.org/package=Rserve), which allows a java based application server to remotely connect through TCP/IP to R installed on an AWS cloud-computing instance. Every Rserve connection uses a separate workspace and working directory, allowing analyses to be easily distributed from the application server to multiple computing nodes as required, thus leveraging the cloud-computing infrastructure to enhance performance and deliver scalable analytics through the platform. While WebMeV’s cloud-based design was chosen to provide scalability, in reality the number of computing nodes allocated to the application can affect performance and the number of compute nodes is limited by our funding to support the project. Because we recognize some users may want to run large dedicated analyses, we provide a WebMeV docker container image (https://hub.docker.com/r/cccb/mev-web/) for users to download and deploy on their own hardware architecture to take advantage of the computing resources to which they have access.

A major challenge in integrating a large number of Bioconductor packages under a single application is management of package specific dependencies and versions. To solve this problem, we implemented an inversion dependency framework (https://cran.r-project.org/web/packages/injectoR/index.html) with R version repository (https://github.com/dfci-cccb/raven) that decouples Bioconductor package dependencies from the global Rserve environment. This framework helps WebMeV to tightly control the dependencies being used by defining clear boundaries around each Bioconductor package; this allows for the inclusion of Bioconductor packages with contradicting dependency requirement during implementation.

### Data Visualization

Another factor that greatly affects application efficiency is data visualization. WebMeV emphasizes bringing genomic data to users through the use of informative and interactive visualizations. WebMeV visualizations were primarily constructed using D3.js (4). D3.js is an open-source JavaScript data visualization library that conforms to SVG, HTML5, and CSS open standards and allows for seamless integration with modern browsers. It also provides software developers the flexibility to manipulate any part of the document object model (DOM) such that one can design with little restriction and so can easily produce dynamic and interactive genomic data visualization.

## Features

### Accessing Public Data Sets

While WebMeV allows users to upload their own data for analysis, it was also designed to provide intuitive access to and analysis of large, publicly available data sets, allowing exploration of these data sets and the development of new hypotheses.

One of the most significant advantages of cloud based application systems is that they can “bring the analytics to the data,” alleviating the need for users to acquire and upload data to an application for analysis. WebMeV was developed to provide two routes for deploying and accessing data on the cloud. Since the largest repository of gene expression data is the Gene Expression Omibus (GEO) (5), we used the NCBI E-utilities API to access data directly from GEO and to import normalized gene expression and sample annotation data into WebMeV. We also aggregated and curated data from the TCGA, the largest public source of cancer genomic data in cancer, including both RNA-seq and microarray gene expression data, both of which are stored with annotation in the WebMeV AWS S3 bucket. There are other large expression data sets that users have requested we add, including GTEx (2), which will be included in a future release.

### Analysis Methods

Because there is a significant collection of gene expression analysis tools in Biocondcutor, the Rserve package integration into WebMeV allowed us to substantially decrease the time and effort associated with deploying new tools. Modularizing both client-end data visualization and data analysis also drastically reduce implementation time. For the initial deployment of WebMeV, we concentrated on incorporating gene expression analysis methods for both microarray and RNA-seq data. The RNA-seq methods include a variety of tools for normalization and detection of differentially expressed genes (LIMMA/VOOM (6), edgeR (7), DESeq (8)), for gene set enrichment analysis (topGO, ReactomePA (9)), and tools for survival analysis in a framework that delivers concise, interactive graphical representations of the results [**Figure 1**]. In addition to gene expression data analysis, the WebMeV interactive data visualization capability enables users to formulate data-driven sample groupings by explore unsupervised clustering (hierarchical clustering, k-nearest neighbor clustering) and principle component analysis (PCA) results. Users can interactively select samples groups for additional analysis or comparison using results such as hierarchical clustering tree branches and PCA distribution patterns.

**Figure 1.**
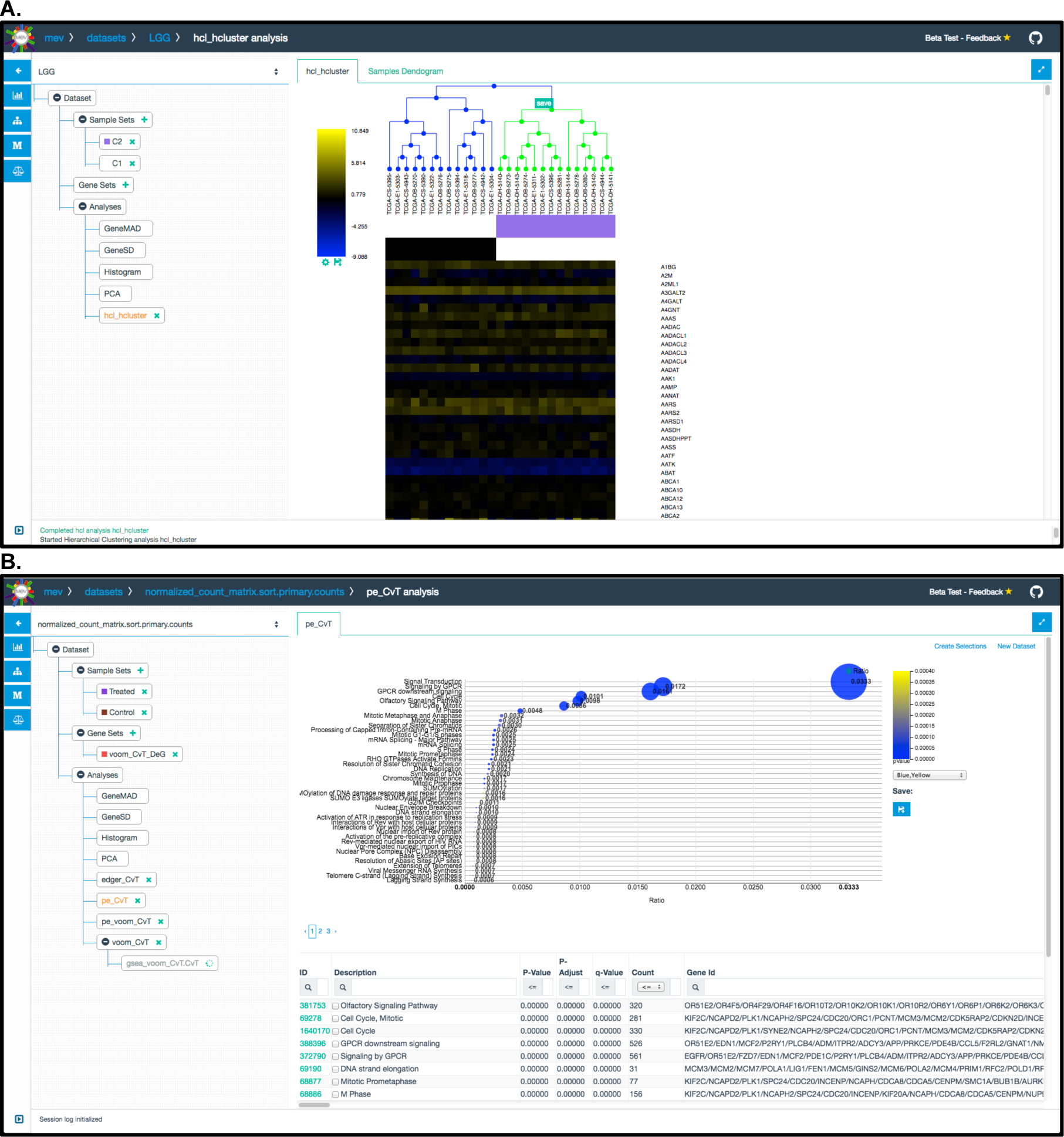
Screenshots of WebMeV showing (A) a simple hierarchical clustering, and (B) a bubble plot summarizing the results of a gene set enrichment analysis.

### Cohort Selection

The basis for most experimental or computational analyses is the identification of distinct cohorts and the comparison of those cohorts to identify differences. WebMeV was designed to allow users to upload data and test specific hypotheses by defining and comparing cohorts. For example, one could use the TCGA data to ask which genes are differentially expressed between HER2+ and ER+ breast cancer patients? What are the biological pathways that appear active in early onset prostate cancer patients compared to those who develop the disease later? Answering these questions requires the construction of complex composite queries on high-dimensional clinical attributes to stratify patient samples into specific cohorts that can then be compared to identify significant differences.

To enable such queries on the TCGA, we implemented a cohort selector in WebMeV using OpenRefine (http://openrefine.org). OpenRefine is an open source application for data cleaning and transformation. OpenRefine has an interface that is similar to a spreadsheet application but behaves like a database in that it allows complex queries and operations on the data. We adapted OpenRefine to support multiple clinical attribute selection methods such as regular expression, numerical range, and categorical facets, to allow users easily visualize attribute distribution across cohort for selection. Users can aggregate multiple selections into a composite criterion for complex cohort stratification and perform set operations on defined cohorts to reduce manual curation effort.

### Example Use Case 1

WebMev was designed for use in a variety of different scenarios. As a first example, consider a simple experiment involving measuring gene expression using RNA-Seq to compare gene expression in three treated samples to that in a set of three control samples. We will assume that the researcher starts with the typical data produced by RNA-Seq experiment : a tab-separated gene-level raw count matrix file (a file containing a set of genes and their RNA-Seq read counts in each sample, with tabs between column entries). The researcher can upload the file to WebMeV and first normalize across all samples. The normalized gene expression distribution can be explored using PCA and experimental and control groups to be contrasted can be defined based on the groupings visualized in the PCA plots. The researcher can also perform unsupervised hierarchical clustering to define sample groups. Or simply separate samples into groups based on treatment status. The groups defined by the user can be compared to identify differentially expressed genes, using a method such as VOOM (6). The differentially expressed genes can then be further filtered and saved as gene list for pathway enrichment analysis. A video showing a step-by-step walkthrough of this example is available at https://youtu.be/6aK4t2vXcg4.

### Example Use Case 2

WebMeV also allows users to download and analyze public data sets such as those from TCGA. Consider a researcher is interested in comparing gene expression between breast cancer patients differing in HER2, PR, and ER status using all the available breast invasive carcinoma (TCGA-BRCA) RNA-Seq data. The researcher can launch WebMeV, select TCGA data, identify and import the full level 3 TCGA-BRCA data set. Data and metadata from TCGA are uploaded and the researcher can use the text facet function in the Sample Set Selector to stratify the cohort based on hormone receptor status. Once individual sub-cohorts are defined, the researcher can use the Sample Sets operation to construct sub-cohorts with composite criteria such as patients with triple negative status. Cohorts or sub-cohorts can be compared to identify differentially expressed genes and to perform a meta-analysis. The gene expression pattern of each sample group can be visualized and explored, including using PCA. A video demonstrating this example is available at https://youtu.be/W48Zhouzdr4.

## Conclusions

In genomics and computational biology. there is too often a divide between those who generate the data and those who analyze it. A physician scientist treating ovarian cancer patients will approach the analysis of ovarian cancer gene expression data with a very different mindset than a computational biologist or biostatistician and, if able to follow his instincts through hands-on analysis, is very likely to reach different insights that if working through a quantitative scientist as an intermediary. Our goal in developing WebMeV has been to provide such a resource for the community with a particular emphasis on assuring that large-scale RNA-seq and other genomic data sets are accessible to the wide range of potential users, In particular, we want to assure that basic, clinical, and translational scientists have access to large public data sets such as those from the TCGA and other public projects in a tool that allows them to explore the data and develop and test hypotheses—and to extend those explorations to their own data. While we recognize the importance of having quantitative scientists, trained in computational biology, bioinformatic, or biostatistics, involved in analyzing data, scientists who understand cancer has unique intuition that they should be enabled to follow in searching to make new discoveries.

## Acknowledgements

The authors would like to thank Alex Holman PhD, Brian Lawney PhD for thoughtful discussions and initial testing.

## References

1. Cancer Genome Atlas Research Network, Weinstein JN, Collisson EA, Mills GB, Shaw KRM, Ozenberger BA, et al. The Cancer Genome Atlas Pan-Cancer analysis project. Nat Genet. Nature Research; 2013;45:1113–20.

2. GTEx Consortium. Human genomics. The Genotype-Tissue Expression (GTEx) pilot analysis: multitissue gene regulation in humans. Science. 2015;348:648–60.

3. Gentleman RC, Carey VJ, Bates DM, Ben Bolstad, Dettling M, Dudoit S, et al. Bioconductor: open software development for computational biology and bioinformatics. Genome Biology. BioMed Central; 2004;5:1.

4. Cottam JA, Lumsdaine A, Wang P. Abstract rendering: out-of-core rendering for information visualization. Wong PC, Kao DL, Hao MC, Chen C, editors. IS&T/SPIE Electronic Imaging. International Society for Optics and Photonics; 2013; 9017:90170K–90170K–13.

5. Barrett T, Wilhite SE, Ledoux P, Evangelista C, Kim IF, Tomashevsky M, et al. NCBI GEO: archive for functional genomics data sets–update. Nucleic Acids Research. Oxford University Press; 2013;41:D991–5.

6. Ritchie ME, Phipson B, Wu D, Hu Y, Law CW, Shi W, et al. limma powers differential expression analyses for RNA-sequencing and microarray studies. Nucleic Acids Research. Oxford University Press; 2015;43:e47–7.

7. Robinson MD, McCarthy DJ, Smyth GK. edgeR: a Bioconductor package for differential expression analysis of digital gene expression data. Bioinformatics. Oxford University Press; 2010;26:139–40.

8. Anders S, Huber W. Differential expression analysis for sequence count data. Genome Biology. BioMed Central Ltd; 2010;11:R106.

9. Yu G, He Q-Y. ReactomePA: an R/Bioconductor package for reactome pathway analysis and visualization. Mol Biosyst. The Royal Society of Chemistry; 2016;12:477–9.

